# Metal-independent effects of calprotectin on cocultures of *Pseudomonas aeruginosa* and *Staphylococcus aureus* involve alkylquinolone production

**DOI:** 10.64898/2026.04.08.717160

**Authors:** Wei H. Lee, Noah H. Tobin, Amanda G. Oglesby, Elizabeth M. Nolan

## Abstract

The current working model of the innate immune protein calprotectin (CP) focuses on its metal-sequestering activity, which contributes to host defense against infection. Recently, CP was reported to enhance the survival of *Staphylococcus aureus* in coculture with *Pseudomonas aeruginosa* in a metal-independent manner. This prior work indicated that the CP protein scaffold, even in the absence of its metal-binding sites, possesses activities that impact interspecies dynamics between these bacterial pathogens. In this study, we employ ΔΔ, a CP variant lacking both functional metal-binding sites, to assess the responses of each pathogen to the CP protein scaffold in monoculture and coculture. Using dual-species transcriptomics, we report that ΔΔ treatment induced gene expression changes indicative of cell envelope modifications for both *P. aeruginosa* and *S. aureus* during coculture. The presence of the CP protein scaffold also attenuated the production of the quorum sensing molecule C_4_-homoserine lactone and the anti-staphylococcal alkylquinolone (AQ) metabolite 2-heptyl-4-hydroxyquinoline N-oxide. Cocultures with *S. aureus* and *P. aeruginosa* mutants defective in AQ biosynthesis demonstrated that AQ production was required for ΔΔ to impact expression of membrane remodeling genes in both species during coculture. Furthermore, we showed that in the absence of AQ production, the effect of CP on *S. aureus* in coculture resembled that of Fe depletion. Collectively, our findings demonstrate that the functional versatility of CP extends beyond multi-metal sequestration and that its intertwined metal-dependent and -independent activities have important consequences for bacterial physiology and polymicrobial interactions.

**IMPORTANCE:** Recent studies of the innate immune protein calprotectin (CP), which is known for its metal-sequestering ability and contributions to nutritional immunity, have uncovered that the protein also exerts metal-independent activities on bacterial pathogens. In this work, we investigate the metal-independent effects of CP on the interspecies dynamics of *Pseudomonas aeruginosa* and *Staphylococcus aureus*, two high-priority pathogens that co-colonize various polymicrobial infection sites. We report that the ability of the CP protein scaffold to attenuate the anti-staphylococcal activity of *P. aeruginosa* results from perturbed quorum sensing and reduced production of alkylquinolone (AQ) metabolites. We further show that pseudomonal AQs contribute to cell envelope remodeling responses exhibited by both pathogens in the presence of the CP protein scaffold. These results afford an updated working model wherein both canonical metal-dependent and noncanonical metal-independent activities of CP elicit physiological changes in both pathogens, resulting in perturbed coculture dynamics.

## INTRODUCTION

The innate immune protein calprotectin (CP, S100A8/S100A9 heterooligomer) defends the host against microbial infection by sequestering multiple nutrient metal ions that are required for the survival and proliferation of pathogens (1–3). This process occurs as part of a host-defense strategy termed nutritional immunity (3–5). CP possesses two metal-binding sites, a His_3_Asp site that binds Zn(II) with high affinity, and a His_6_ site that binds multiple divalent transition metal ions with high affinity (2, 6). To date, studies examining the interplay between CP and microbial pathogens have primarily focused on the battle for metal nutrients at the host–pathogen interface. These efforts have shown that CP exerts antimicrobial activity against a wide spectrum of microbial pathogens (3, 5, 7), and this activity has been broadly attributed to the ability of CP to starve these pathogens of essential metal nutrients (1, 6, 8, 9). *Pseudomonas aeruginosa* and *Staphylococcus aureus* are two prominent bacterial pathogens that have been extensively studied in the context of CP and nutritional immunity (6, 10–12). These pathogens co-colonize multiple polymicrobial infection sites, including chronic wounds and the lungs of individuals with cystic fibrosis (CF). Recent studies have examined how *P. aeruginosa* and *S. aureus* respond to the presence of CP in monocultures, revealing how these bacterial pathogens use multiple strategies to cope with CP-mediated metal limitation (11, 13–19). In addition, several investigations described below have considered how CP affects these two pathogens in coculture (14, 17, 18). Despite major advances from this combined work, our understanding of how CP affects the physiologies of *P. aeruginosa* and *S. aureus* remains incomplete, especially in the context of interspecies interactions that are relevant to many infection states, necessitating further investigations to elucidate how CP affects the these pathogens.

Interactions between *P. aeruginosa* and *S. aureus* are often antagonistic, and coculture of both species frequently results in killing of *S. aureus* by *P. aeruginosa* (20, 21). Importantly, two independent studies investigating how nutritional immunity impacts this polymicrobial interaction reported that the presence of CP promoted the survival of *S. aureus* in coculture with *P. aeruginosa* (14, 17). One of these studies showed that CP treatment of *P. aeruginosa* monocultures also decreased the levels of alkylquinolones (AQs) (14), pseudomonal metabolites that possess anti-staphylococcal activity (20, 22, 23). Subsequently, we investigated the consequences of CP treatment on *P. aeruginosa* and *S. aureus* cocultures using dual-species transcriptomics and mass spectrometry (MS). We found that CP treatment perturbed chorismate flux, attenuated quorum sensing (QS), and reduced AQ production by *P. aeruginosa* during coculture (18). In addition, our work showed that CP treatment elicited gene expression responses indicative of cell envelope remodeling from both pathogens in coculture (17, 18). These findings led to a working model describing the impact of CP on coculture dynamics between *P. aeruginosa* and *S. aureus* (**Figure 1**). In this model, CP dampens pseudomonal QS and decreases AQ production, particularly levels of the AQ N-oxides (AQNOs) that have been most closely correlated with anti-staphylococcal activity in multiple reports (18, 20, 24).

**Figure 1.**
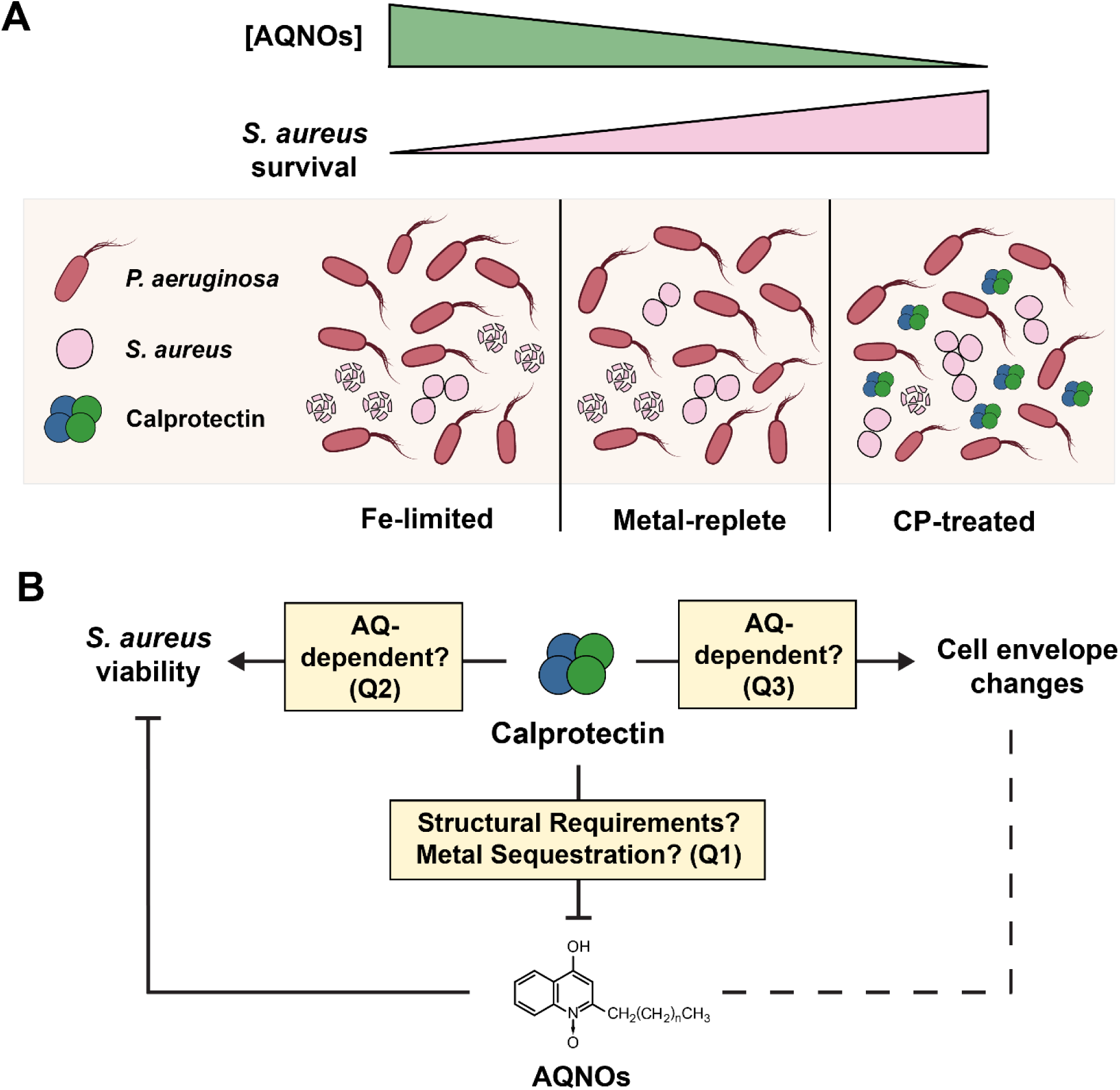
The opposing effects of CP treatment and Fe limitation on coculture dynamics between *P. aeruginosa* and *S. aureus* (**A**), and outstanding questions for the current working model describing the impact of CP on coculture dynamics (**B**). Which structural features of CP are required for the observed activity on gene expression and AQ production in *P. aeruginosa* / *S. aureus* cocultures (**Q1**)? Is *S. aureus* protection driven by CP-induced cell envelope remodeling, perturbed AQ production, or other processes (**Q2**)? Are cell envelope remodeling responses observed in both pathogens caused by altered AQ production (**Q3**)? Panel A was reproduced from ref (18).

Curiously, our studies also showed that CP treatment and Fe depletion exert opposing effects on *S. aureus* viability during coculture with *P. aeruginosa* (**Figure 1A**). While CP treatment reduced AQ production and protected *S. aureus*, Fe depletion enhanced anti-staphylococcal activity by increasing AQ production (17, 18, 23). These results presented an apparent dichotomy in the manner by which CP affects interactions of these two pathogens. Towards addressing this dichotomy, we examined the effect of ΔΔ, a site variant of CP lacking both the His_3_Asp and His_6_ metal-binding sites that is unable to sequester transition metal ions, on coculture dynamics (17). We found that CP-mediated protection of *S. aureus* in coculture with *P. aeruginosa* occurs without metal sequestration, uncovering an underappreciated metal-independent activity of this protein. Additionally, ΔΔ affected expression of genes indicative of cell envelope remodeling in both species. Together, these observations demonstrated that other aspects of CP beyond its metal-withholding ability modulate interspecies interactions.

Considering these findings in the context of the existing working model, several outstanding questions remain to be addressed (**Figue 1B**). For example, further studies are necessary to determine how the CP protein scaffold contributes to changes in gene expression and AQ production during coculture (**Figure 1B, Q1**). Additionally, it is unclear whether the protective effect of CP on *S. aureus* in coculture is attributable to changes in AQ production, AQ-independent processes, or a combination of such factors (**Figure 1B**, **Q2**). Moreover, the current model does not address how altered AQ production contributes to cell envelope remodeling responses that are elicited by CP during coculture of these two pathogens (**Figure 1B**, **Q3**) (17, 18). Insights gained from addressing these questions will advance the understanding of how CP, independent of metal sequestration, modulates microbial physiologies that underlie interspecies interactions.

In the current study, we investigated whether the CP protein scaffold can recapitulate the transcriptional responses elicited by CP by performing dual-species RNA-seq of *P. aeruginosa* and *S. aureus* cocultures treated with and without ΔΔ. Through analysis of these data and our prior CP treatment dataset (18), we delineated the effects of CP-mediated metal sequestration and the CP protein scaffold on the gene expression profiles of each species in coculture. Using quantitative MS, we demonstrated that ΔΔ treatment decreases levels of the QS molecule *N*-butanoyl-L-homoserine lactone (C_4_-HSL) during early growth and suppresses production of the anti-staphylococcal metabolite 2-heptyl-4-hydroxyquinoline N-oxide (HQNO) (25–27). Furthermore, we employed genetic mutants of *P. aeruginosa* that lack one or more of the *pqs* biosynthetic genes to show that, while the AQNOs are major drivers of *P. aeruginosa* / *S. aureus* coculture outcomes, additional AQ metabolites contribute to coculture dynamics. Our study also indicates that the ability of CP and ΔΔ to alter gene expression changes related to cell envelope remodeling of *P. aeruginosa* and *S. aureus* relies on perturbed AQ production. Moreover, in the absence of AQ production by *P. aeruginosa*, the impact of CP on *S. aureus* viability and transcriptional responses in coculture largely resembles that of Fe depletion, whereas ΔΔ treatment exerts negligible effects on *S. aureus*. Overall, these findings demonstrate how the metal-dependent and -independent activities of CP exert intertwined effects on bacterial physiologies and interactions.

## RESULTS

### The presence of the CP protein scaffold perturbs *S. aureus* and *P. aeruginosa* gene expression during coculture

The RNA-seq experimental design followed our recent transcriptomics studies of *P. aeruginosa* and *S. aureus* cocultures (18). In brief, *P. aeruginosa* PA14 and *S. aureus* USA300 JE2 were grown in metal-replete chemically defined medium (CDM), a growth medium prepared from trace metals basis reagents and supplemented with defined concentrations of Ca, Mn, Fe, Ni, Cu, and Zn immediately prior to use (8, 17, 28). We purified ΔΔ following previously established procedures (17, 29); like CP, this variant exhibits the expected secondary structure and Ca(II)-dependent self-association to form S100A8/S100A9 heterotetramers (7, 17). To investigate how the CP protein scaffold affects *P. aeruginosa* / *S. aureus* interspecies dynamics, we performed dual-species RNA-seq on cocultures of these two pathogens grown in metal-replete CDM with or without 20 *µ*M ΔΔ. We also sequenced transcripts from monocultures of *P. aeruginosa* and *S. aureus* treated with or without ΔΔ (20 *µ*M) to identify gene expression changes that were common to both coculture and monoculture, or unique to either culture type. The presence of ΔΔ did not significantly affect the viability of *P. aeruginosa* or *S. aureus* in coculture or monoculture at the 6 h timepoint (**Figure S1**) (17). RNA-seq was performed at the 6 h timepoint based on considerations identified in prior work and to allow for comparison of the current ΔΔ datasest with our prior CP dataset (17, 18). For differential expression analysis, we considered genes with expression levels that differed by at least 1 log_2_(fold change) (LFC) relative to untreated cultures to be differentially expressed (DE). The proportion of genes that were DE in each condition are reported in **Table S1**, and the key statistics of the RNA-seq dataset are summarized in **Table S2**.

Differential expression analysis of the sequenced reads for *P. aeruginosa* revealed that less than 2% of *P. aeruginosa* genes were DE in both monocultures and cocultures treated with ΔΔ (**Figures 2A**, **S2A** and **Table S1**), which was much smaller than the proportion of DE genes (approximately 25%) detected in CP-treated monocultures and cocultures (18). We found that ΔΔ treatment resulted in no statistically significant transcriptional changes for *S. aureus* in monoculture (**Figure S2B** and **Table S1**) and only about 6% of *S. aureus* genes in coculture were DE in response to ΔΔ treatment (**Figures 2B**, **S2B** and **Table S1**). This comparison indicated that the response of *S. aureus* to the CP protein scaffold was largely dependent on the presence of *P. aeruginosa*. The overall proportion of DE genes for *S. aureus* in ΔΔ-treated cultures was also decreased as compared to CP treatment (monoculture: ∼7%, coculture: ∼40%) (18). We performed gene set enrichment analysis of the DE *P. aeruginosa* genes (**Figure S4**) and attempted to perform similar functional enrichment analyses for the *S. aureus* genes. However, these efforts did not yield any statistically significant gene clusters for the latter, which we believe is in part due to the small number of *S. aureus* genes fulfilling the LFC threshold as well as limited Gene Ontology (GO) annotations available (18). The full lists of DE *P. aeruginosa* and *S. aureus* genes for each condition are tabulated as a **Supplementary File** (**Tables SF1** and **SF2**).

**Figure 2.**
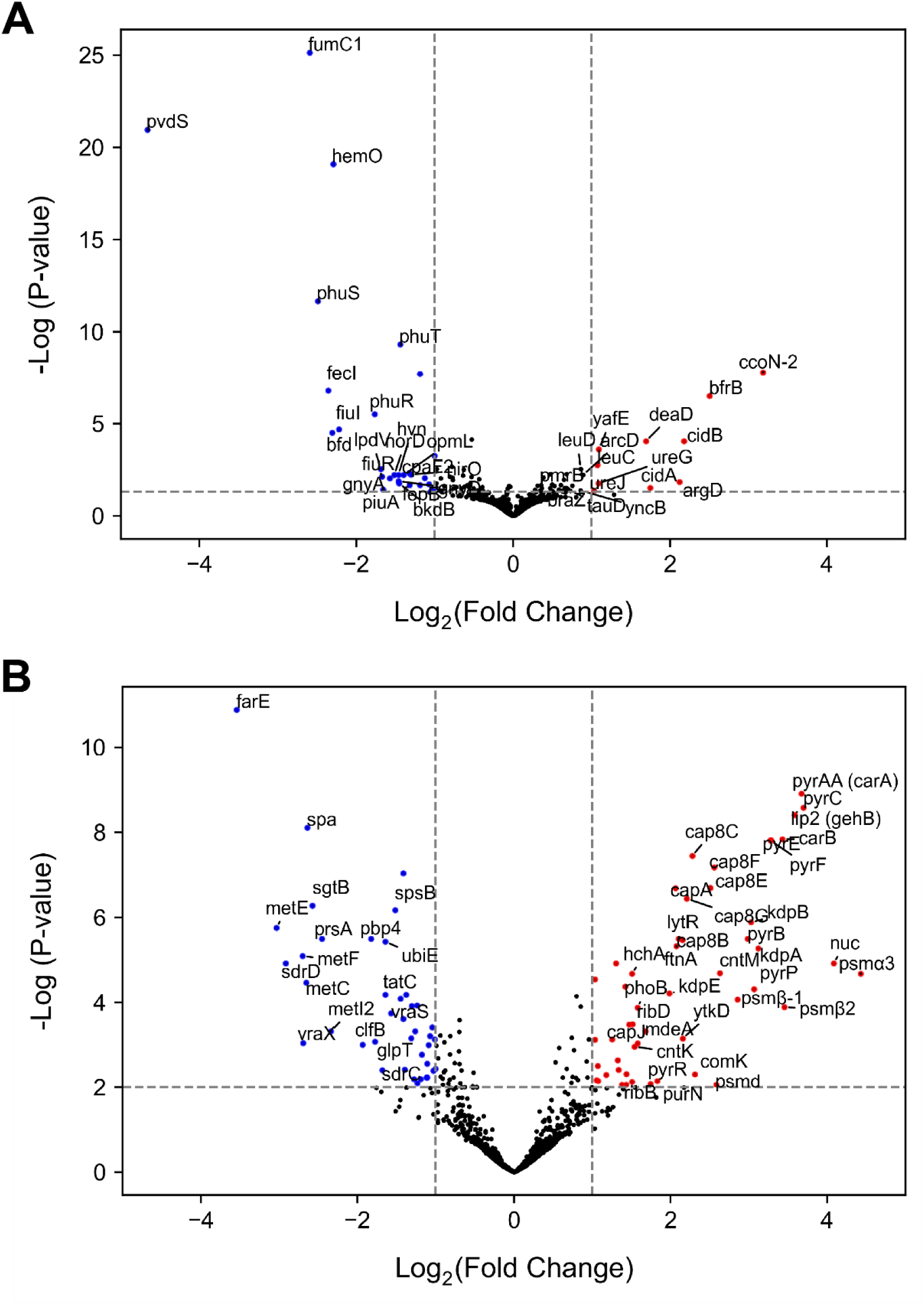
Volcano plot of annotated *P. aeruginosa* (**A**) and *S. aureus* (**B**) genes DE in response to ΔΔ treatment in coculture. Genes which were DE with a P-value < 0.01 are colored; red denotes upregulated (LFC > 1) and blue denotes downregulated (LFC<1).

Consistent with the inability of ΔΔ to sequester metal ions, metal-starvation responses such as the upregulation of machinery involved in siderophore biosynthesis, metal acquisition, and stress responses were not observed for *P. aeruginosa* in coculture (**Figure 2A**). In the same vein, the presence of ΔΔ did not result in the induction of hallmark Fe-starvation responses such as siderophore biosynthesis and heme uptake for *S. aureus* in coculture (**Figure 2B**). Unexpectedly, we observed that ΔΔ treatment resulted in gene expression changes for both pathogens indicative of altered cellular metal requirements or metal homeostasis (**Figures 2A** and **2B**, see following subsection). Moreover, the presence of ΔΔ resulted in gene expression changes indicative of cell envelope modifications for both *P. aeruginosa* and *S. aureus* in coculture (**Figures 2A** and **2B**). Each of these findings are described in more detail below.

#### Treatment with the CP protein scaffold affects the expression of genes associated with metal homeostasis for *P. aeruginosa* and *S. aureus* in coculture

For *P. aeruginosa*, ΔΔ treatment resulted in significant downregulation of multiple genes associated with Fe acquisition and utilization (**Figures 2A** and **S3A**); these genes comprised the six largest clusters of GO terms detected by gene set enrichment analysis for *P. aeruginosa* in ΔΔ-treated cocultures (**Figure S4**). The downregulated genes included those encoding pyochelin (PCH) biosynthetic machinery (30), the sigma factor PvdS, which regulates pyoverdine biosynthesis (31, 32), heme uptake and utilization machinery (Phu and HemO) (33, 34), and the Fe-S cluster assembly regulator IscR (∼0.9 LFC, just below the 1.0 LFC cut-off) (**Figure S3A**) (35). The downregulation of *pvdS* was further demonstrated using real-time PCR (**Figure S3B**). The presence of ΔΔ also resulted in the downregulation of genes encoding xenosiderophore transport systems, including *fecI*, *fiuIR*, *femIR*, *pirA* and *piuA* (36–38), the Fe-responsive *PA4468*–*PA4471* operon (39, 40), which includes the genes for the Fe-independent fumarase analog FumC1 (40, 41), and *sodM*, which enodes the cambialistic Mn/Fe-cofactored superoxide dismutase SodM (**Figure S3A**). Furthermore, upregulation of the genes for bacterioferritin (*bfrB*) and downregulation of its associated ferredoxin (*bfd*) suggested that *P. aeruginosa* in ΔΔ-treated cocultures was sensing higher cytosolic Fe levels (**Figure S3A**) (43–47). We observed that ΔΔ treatment resulted in a relatively modest effect on Fe-starvation responses in *P. aeruginosa* monocultures that was restricted to decreased expression of genes for the PCH biosynthetic machinery (**Figure S3A**). For *S. aureus*, ΔΔ treatment resulted in upregulated expression of the ferrous iron transporter gene *feoB* and genes encoding two ferritin family proteins, FtnA and Dps (**Figure S5**). ΔΔ treatment of *S. aureus* in coculture also resulted in the upregulation of *cntMLK*, genes encoding the biosynthesis of the nicotianamine-like metallophore staphylopine (**Figure S5**) (48, 49). Overall, these results suggested that ΔΔ affects metal homeostasis of both pathogens during coculture, an unexpected finding that warrants future investigation.

#### The presence of the CP protein scaffold induces transcriptional responses indicative of cell envelope modifications in *P. aeruginosa* and *S. aureus* during coculture

Our transcriptomics further showed that ΔΔ treatment of *P. aeruginosa* / *S. aureus* cocultures resulted in the upregulation of *P. aeruginosa* genes associated with cell envelope modifications. These upregulated genes included *cidAB*, the products of which modulate the release of extracellular DNA (eDNA) in *P. aeruginosa* (50–52), and *arnT* (∼0.7 LFC, falling just under the LFC threshold), which is involved in lipid A modification (**Figure 2B**) (53, 54). ΔΔ treatment also resulted in upregulation of the *pmrAB* two-component system (**Figure 3A**), which contributes to the resistance of *P. aeruginosa* against cationic antimicrobial peptides (55–57). Our prior findings suggested that both CP-mediated Zn(II) sequestration and the CP protein scaffold perturb the composition of the *P. aeruginosa* outer cell membrane and induce cell envelope remodeling responses for *P. aeruginosa* cocultured with *S. aureus* (13, 17, 18). The presence of ΔΔ also led to downregulation of genes associated with the *pprAB* regulon (**Figure 3A**) (by ∼0.6–2.0 LFC), which includes genes encoding CupE fimbriae (58) and is thought to be involved in a distinct, exopolysaccharide-independent mode of *P. aeruginosa* adhesion to surfaces (59). Within the *pprAB* regulon, we identified *flp*, which encodes for a type IVb pilin, as a candidate gene for further validation of our transcriptomics data (60, 61). Using real-time PCR, we observed that ΔΔ treatment resulted in a significant decrease in *flp* transcript levels (**Figure 3B**). Taken together, these findings are overall consistent with gene expression trends observed for *P. aeruginosa* in CP-treated cocultures (13, 17, 18). The results for the *pprAB*-associated regulon (**Figure 3A**) mirrored transcriptional changes observed in CP-treated cocultures (∼2.5–5.5 LFC), although the magnitude of this effect was attenuated under ΔΔ treatment (18, 60). One possibility worth considering is that the presence of the CP protein scaffold may affect surface sensing and adhesion by *P. aeruginosa*, diminishing the expression of type IV pili and inducing cell envelope remodeling responses (62).

**Figure 3.**
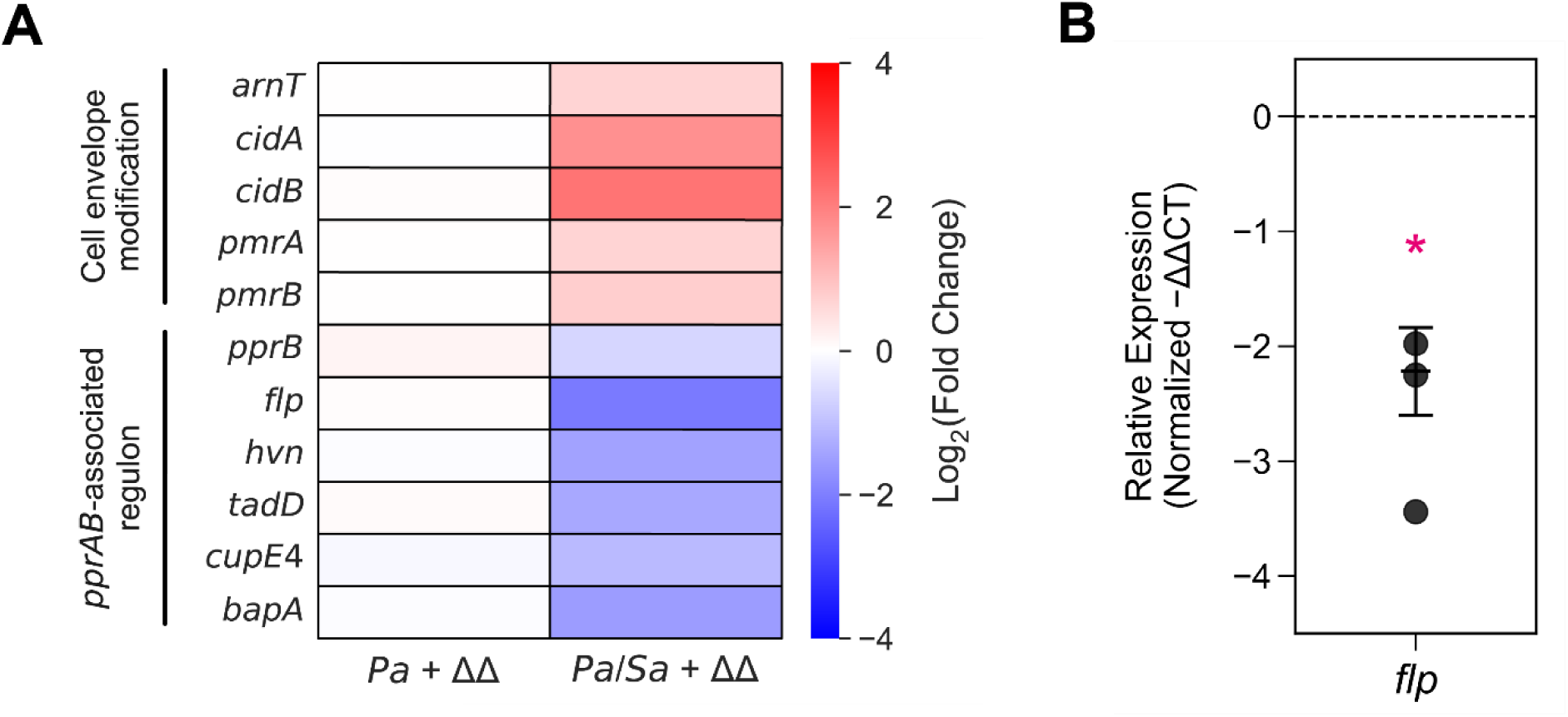
The CP protein scaffold upregulates the expression of genes associated with cell envelope modifications and downregulates the *pprAB*-associated regulon in *P. aeruginosa* cocultured with *S. aureus*. DE heatmap of *P. aeruginosa* genes associated with cell envelope modifications and the *pprAB*-associated regulon (**A**), and the change in expression for the type IV pili gene *flp* in response to ΔΔ treatment (**B**). *Pa* indicates *P. aeruginosa* monoculture and *Pa/Sa* indicates the coculture. For real-time PCR, transcript levels were normalized to the 16S housekeeping gene for *P. aeruginosa*, and the normalized Log_2_(Fold Change) is presented. For comparison with the untreated culture condition, \**P* < 0.05.

Prior studies examining the impact of the CP protein scaffold also revealed that ΔΔ induces cell envelope modifications in *S. aureus* when cocultured with *P. aeruginosa* (17). In line with these observations, the presence of ΔΔ increased the expression of genes involved in the production of type 8 capsular polysaccharide and staphyloxanthin (**Figure 4**) (63–65). ΔΔ treatment also downregulated the expression of several genes associated with cell envelope modifications, including (i) genes encoding the peptidoglycan glycosyltransferase SgtB, which cross-links peptidoglycans; (ii) the serine-aspartate repeat proteins, which facilitate *S. aureus* surface adhesion and biofilm formation; and (iii) the twin-arginine translocation system (66–69). Additionally, we observed that ΔΔ treatment resulted in decreased expression of genes encoding the LytSR two-component system (**Figure 4**), a regulator of murein hydrolase activity in *S. aureus* that is essential for membrane remodeling and cellular growth (70, 71). We also detected upregulation of genes encoding the antiholin-like proteins LrgAB and the competence protein ComK, as well as genes coding for competence pili (**Figure 4**), which play a role in binding and uptake of DNA (17, 50, 72–75). Furthermore, ΔΔ treatment led to the DE of several genes associated with membrane transport. Genes encoding the ATP-dependent K^+^ transporter Kdp, the phosphonate import system Phn, and the multidrug efflux transporter MdeA were upregulated in the presence of ΔΔ, whereas genes encoding the Na^+^/H^+^ antiporter Mnh1 and the antimicrobial fatty acid resistance efflux pump FarE were downregulated (**Figure S6**) (76–81). These findings indicated that, during coculture with *P. aeruginosa,* the CP protein scaffold induces *S. aureus* responses indicative of altered eDNA levels, osmotic stress, and/or perturbed intracellular pH regulation.

**Figure 4.**
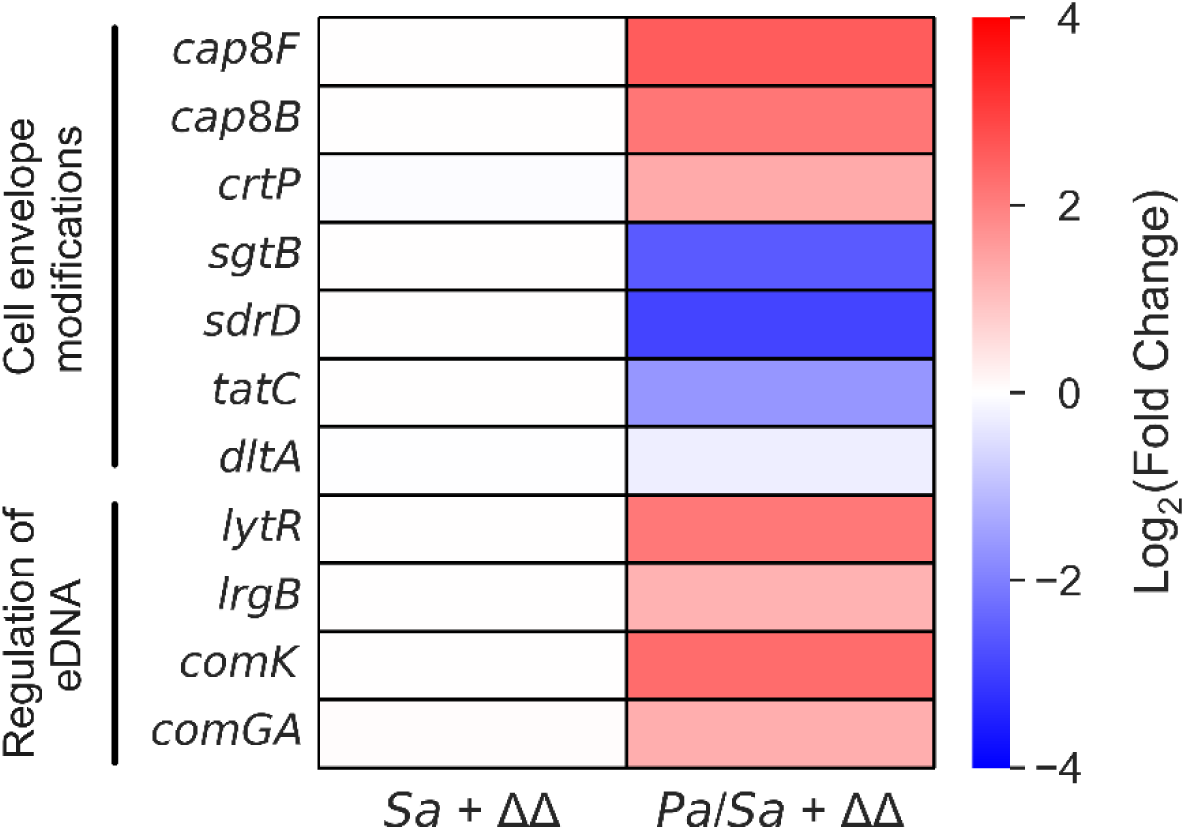
The CP protein scaffold induces transcriptional responses indicative of cell envelope modifications and eDNA remodeling in *S. aureus* cocultured with *P. aeruginosa*. DE heatmap of *S. aureus* genes associated with cell envelope modifications and the regulation of eDNA. *Sa* indicates *S. aureus* monoculture and *Pa/Sa* indicates the coculture.

### The CP protein scaffold perturbs QS in *P. aeruginosa* cocultured with *S. aureus* by decreasing C_4_-HSL production during early growth

Prior studies investigating how CP modulates interspecies dynamics between *P. aeruginosa* and *S. aureus* in coculture indicated that attenuated QS in *P. aeruginosa* and decreased production of pseudomonal antimicrobials in the presence of CP contribute to increased *S. aureus* survival (18). Because ΔΔ was previously reported to confer similar protection to *S. aureus* cocultured with *P. aeruginosa*, we reasoned that the protective effect of CP was most likely metal-independent and hypothesized that the CP protein scaffold would recapitulate the effects of CP on pseudomonal QS in coculture (17, 18). Two of the main signaling molecules involved in *P. aeruginosa* QS are the acylhomoserine lactone (HSL) metabolites 3-oxo-C_12_-HSL and C_4_-HSL, which autoregulate expression of the LasIR and RhlIR systems, respectively (26, 82–84). LasIR and RhlIR are, in turn, part of a hierarchy of QS signaling cascades that control the regulation of pseudomonal virulence factors (85–89). Differential expression analysis showed that ΔΔ treatment resulted in slight but statistically significant decreases in expression of the genes encoding the QS-regulated virulence factors LasA (−0.56 LFC) and LasB (−0.92 LFC), though falling short of the LFC threshold. We monitored the effect of ΔΔ treatment on *lasA* expression using real-time PCR and found that the presence of ΔΔ significantly decreased *lasA* expression (**Figure 5A**). Because the expression of *lasA* is controlled by QS (90–92), we suspected that ΔΔ treatment might perturb QS for *P. aeruginosa* cocultured with *S. aureus*. This notion is also supported by prior studies demonstrating that CP treatment decreased levels of C_4_-HSL to a larger extent than Fe depletion in *P. aeruginosa* cocultured with *S. aureus* (18), suggesting that the CP protein scaffold may act to perturb C_4_-HSL production by *P. aeruginosa*.

**Figure 5.**
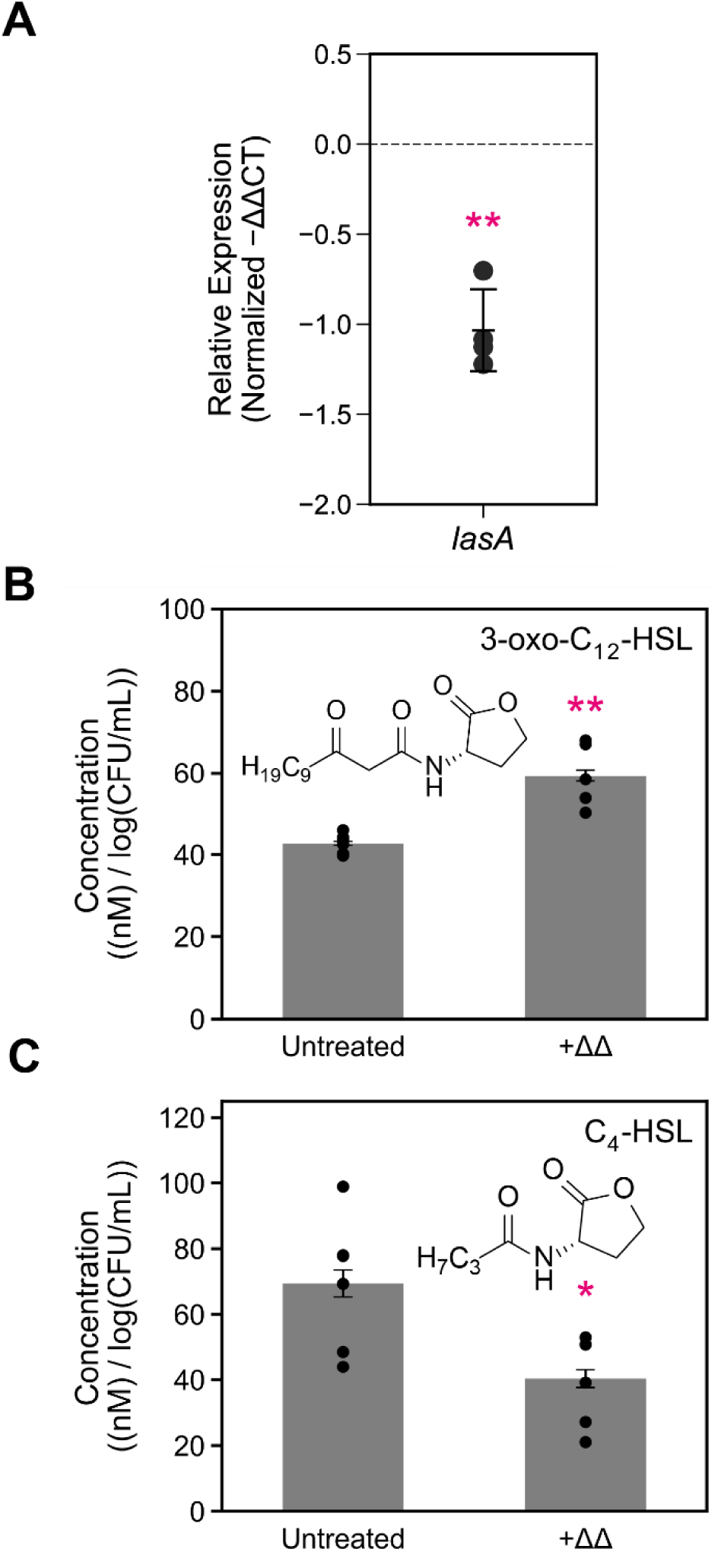
The presence of the CP protein scaffold decreases the expression of *lasA* (**A**), increases levels of 3-oxo-C_12_-HSL (**B**) and decreases levels of C_4_-HSL (**C**) during early growth of *P. aeruginosa* cocultured with *S. aureus*. Aliquots of culture supernatants were collected from cocultures of *P. aeruginosa* and *S. aureus* grown for 6 h in metal-replete CDM ± 20 *µ*M ΔΔ at 37 °C, and processed for quantitative mass spectrometry. Metabolite levels were normalized to *P. aeruginosa* CFUs (n=5, error bars represent S.E.). For comparison with the untreated culture condition, \**P* < 0.05, \*\**P* < 0.01.

Motivated by the overlap between the transcriptional responses detected in response to CP and ΔΔ treatment, we employed triple quadrupole MS to probe the levels of *P. aeruginosa* QS and antimicrobial metabolites (18). To determine whether the presence of ΔΔ perturbed *P. aeruginosa* QS, we quantified levels of 3-oxo-C_12_-HSL and C_4_-HSL in supernatants derived from *P. aeruginosa* / *S. aureus* cocultures treated with or without ΔΔ. These analyses revealed that ΔΔ treatment increased levels of 3-oxo-C_12_-HSL and decreased levels of C_4_-HSL at the 6 h timepoint (**Figures 5B** and **5C**) but did not significantly alter levels of these two QS metabolites at the 11 h timepoint (**Figures S7A** and **S7B**). We also observed that the increased levels of 3-oxo-C_12_-HSL contrasted with the decreased expression of *lasA* at the 6 h timepoint (**Figure 5A** and **5B**). These findings indicated that the CP protein scaffold perturbs *P. aeruginosa* QS during early growth in coculture and suggested that levels of 3-oxo-C_12_-HSL and C_4_-HSL return to untreated levels by the time *P. aeruginosa* reaches stationary phase growth. Altered HSL production was unlikely due to changes in growth rate or phase since the presence of the ΔΔ does not appreciably alter *P. aeruginosa* growth dynamics in coculture and in monoculture (**Figure S1A**) (17).

### The CP protein scaffold reduces HQNO production in *P. aeruginosa* cocultured with *S. aureus*

*P. aeruginosa* produces multiple AQs, some of which participate in QS and others that possess direct anti-staphylococcal activity (25, 93–96). We previously demonstrated that CP-mediated survival of *S. aureus* in coculture with *P. aeruginosa* correlated with decreased production of several AQ metabolites involved in QS [C_7_ AQs 2-heptyl-4(1*H*)-quinolone (HHQ), 2-heptyl-3-hydroxy-4(1*H*)-quinolone (PQS), and C_9_ AQ 2-nonyl-4(1*H*)-quinolone (NHQ)], as well as the anti-staphylococcal AQ HQNO (18). To determine whether ΔΔ elicited similar effects on AQ production as CP, we quantified AQ levels in cocultures treated with ΔΔ by quantitative mass spectrometry.

At the 6 h timepoint, we observed a significant decrease in both HHQ and HQNO upon ΔΔ treatment (**Figure 6A** and **6B**); these decreases were attenuated for HQNO and absent for HHQ at the 11 h timepoint (**Figures S8A** and **S8B**). By contrast, ΔΔ treatment resulted in significantly increased levels of NHQ and C_9_-PQS at the 6 h timepoint (**Figures 6E** and **6G**), though these trends were reversed at the 11 h timepoint (**Figures S8E** and **S8G**). ΔΔ treatment resulted in no significant changes in either PQS or NQNO at either timepoint (**Figures 6C**, **6F**, **S8C** and **S8F**). When considering the impact of each AQ metabolite among the total AQ pools at each timepoint, we noted that HHQ contributed the most to overall changes in the C_7_ congener pool (**Figures 6D** and **S8D**), while both NHQ and C_9_-PQS contributed the most to overall changes in the C_9_ congener pool (**Figures 6H** and **S8H**).

**Figure 6.**
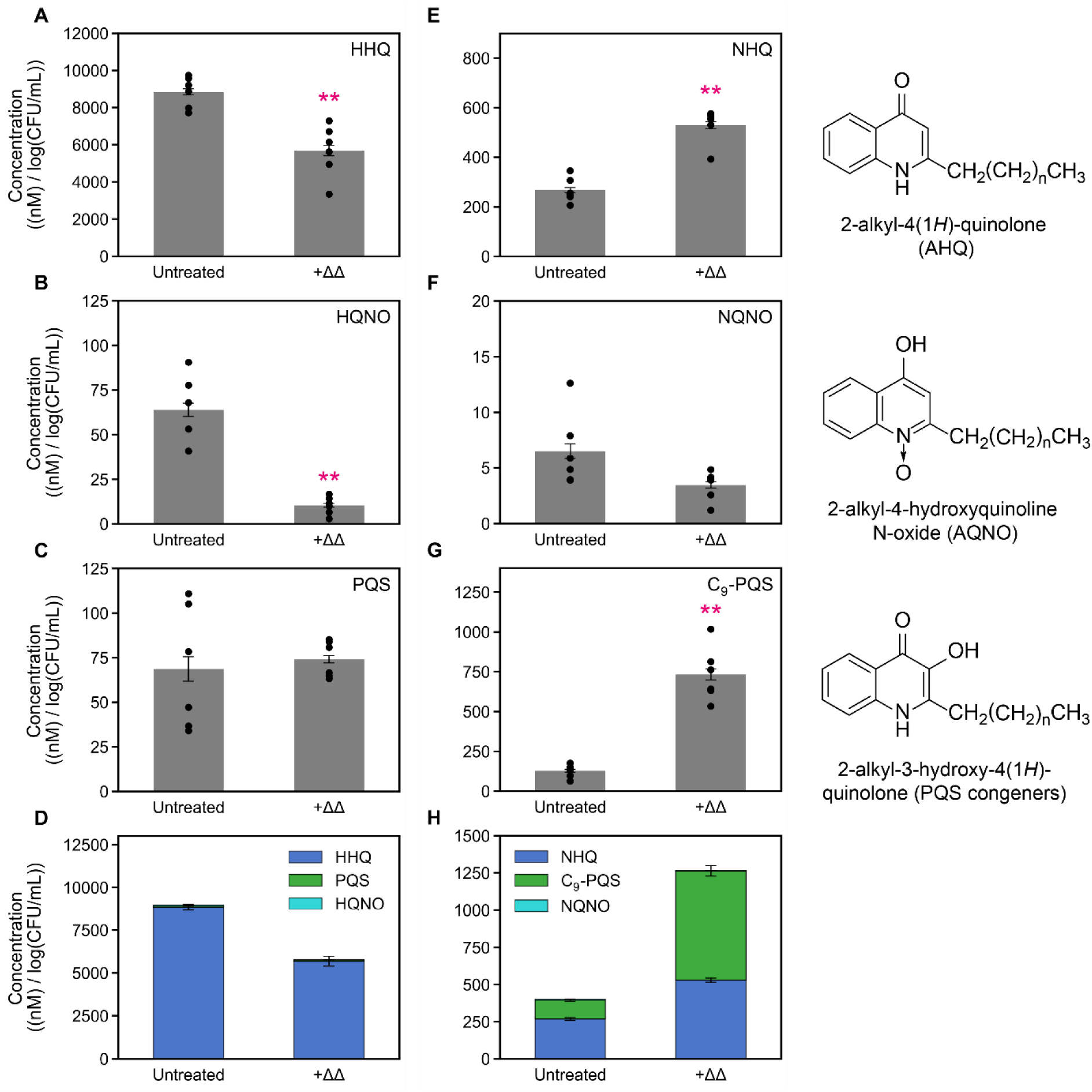
Treatment with ΔΔ decreases the production of C_7_ AQs HHQ and HQNO and increases levels of the C_9_ AQs NHQ and C_9_-PQS during early growth in *P. aeruginosa* / *S. aureus* co-cultures. Treatment with ΔΔ decreases the levels of HHQ (**A**), HQNO (**B**), but not PQS (**C**), resulting in decreased overall levels of C_7_ AQs (**D**). ΔΔ treatment increased the levels of NHQ (**E**), did not significantly affect NQNO levels (**F**), and increased levels of C_9_-PQS (**G**) and the overall C_9_ AQ pool (**H**). For the C_7_ (heptyl) and C_9_ (nonyl) congeners, the alkyl chain length is 7 and 9 carbons long, respectively. Aliquots of culture supernatants were collected from cocultures of *P. aeruginosa* and *S. aureus* grown for 6 h in metal-replete CDM ± 20 *µ*M ΔΔ at 37 °C, and processed for quantitative mass spectrometry. Metabolite levels were normalized to *P. aeruginosa* CFUs (n ≥ 5, error bars represent S.E.). For comparison with the untreated culture condition, \*\**P* < 0.01.

We next considered these changes in the context of prior studies of CP and Fe depletion on AQ production. We noted that changes in overall pools of either the C_7_ or C_9_ AQ congeners did not correlate with changes in *S. aureus* viability across coculture conditions. For example, CP/ΔΔ treatment and Fe depletion resulted in decreased pools of the C_7_ AQ congeners, yet CP/ΔΔ treatment and Fe depletion exerted opposing effects on *S. aureus* viability (**Figures 6D** and **S9**) (17, 18, 20, 23, 24). Instead, we found that changes in HQNO levels inversely correlated with *S. aureus* viability across all studies, as HQNO was reduced upon both ΔΔ and CP treatment and increased upon Fe depletion at both the 6 and 11 timepoints (**Figures 6B**, **S8B**, **S9B** and **S10B**) (14, 18, 24). Combined, these data were consistent with a model in which changes in HQNO production underlie the coculture dynamics of *P. aeruginosa* and *S. aureus* in the presence of CP, ΔΔ, and Fe depletion.

Overall, the MS analyses demonstrated that the presence of the CP protein scaffold, even without functional metal-binding sites, is sufficient to significantly impact AQ production by *P. aeruginosa* in coculture with *S. aureus*. These effects included decreased levels of the potent anti-staphylococcal metabolite HQNO and perturbed levels of the C_9_ AQs NHQ and C_9_-PQS. Taken together, our metabolite analyses suggested that the CP protein scaffold impacts AQ production primarily through perturbation of *P. aeruginosa* QS (**Figure 1B, Q1**). We speculate that the time-dependent effect of ΔΔ treatment on NHQ and C_9_-PQS production could be a response of *P. aeruginosa* to perturbed QS, as both NHQ and C_9_-PQS function as QS metabolites (97–100).

### The effects of CP and ΔΔ on interspecies dynamics in *P. aeruginosa* / *S. aureus* cocultures require AQ production

Data presented here and in our prior work (17, 18) support a model in which CP– and ΔΔ-mediated protection of *S. aureus* during coculture with *P. aeruginosa* results from decreased production of HQNO (**Figure 1A**). To investigate this model further, we employed genetic variants of *P. aeruginosa* lacking one or more components of the PQS biosynthetic pathway. We cocultured *S. aureus* with either the Δ*pqsH* (lacking PQS), Δ*pqsL* (lacking AQNOs), or Δ*pqsA*–*C* (lacking all AQs) deletion mutants constructed in *P. aeruginosa* strain PA14 and analyzed the impact of these mutations on *P. aeruginosa* and *S. aureus* growth during coculture in metal-replete CDM with or without CP or ΔΔ, as well as Fe-depleted CDM (**Figure 7**, **Figures S11** – **S14**).

**Figure 7.**
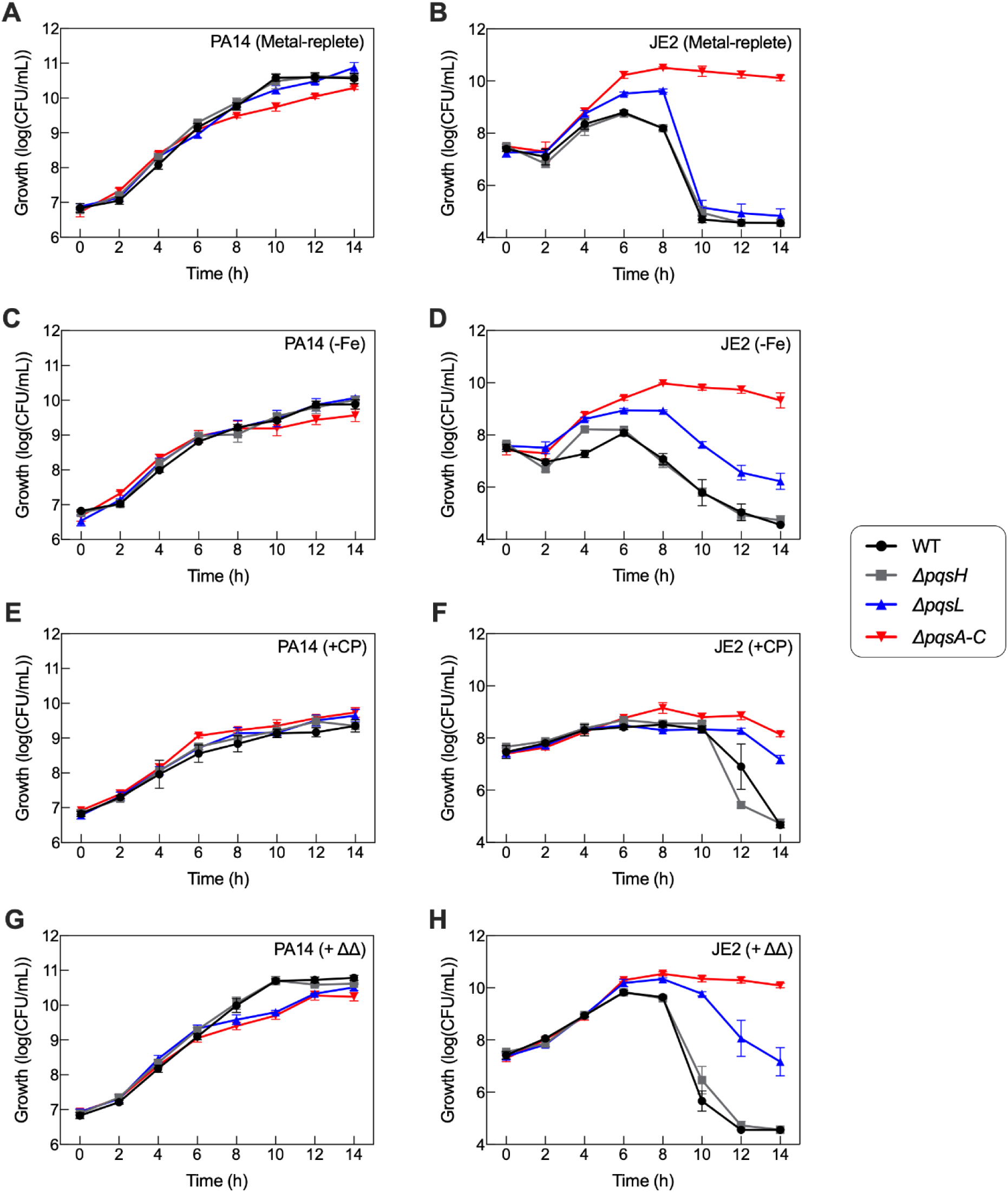
Multiple AQs contribute to the anti-staphylococcal activity of *P. aeruginosa* against *S. aureus* in coculture. Growth dynamics of *P. aeruginosa* wild-type (WT) and Δ*pqs* genetic mutants (**A**, **C, E, G**) and *S. aureus* (**B**, **D, F, H**) in coculture. Cocultures were grown in metal-replete CDM, Fe-depleted CDM, or metal-replete CDM ± 20 *µ*M CP or ΔΔ and incubated at 37 °C (n ≥ 3; error bars represent S.E.). Refer to **Tables S8** and **S9** for significance testing for cell viability at the 8–14 h timepoints.

For cocultures grown in metal-replete CDM, all *P. aeruginosa* strains reached stationary phase around the 10–12 h timepoints, and growth of each *pqs* mutant was comparable to wild-type (**Figure 7A**). CP treatment and Fe depletion of cocultures reduced the growth of all *P. aeruginosa* strains starting at the 6 h timepoint, with a 1–2 log reduction in growth evident by the 14 h timepoint (**Figures S11** – **S14**). In ΔΔ-treated cocultures (**Figure 7G**), viability of the Δ*pqsL* and Δ*pqsA*–*C* mutants was reduced by ∼2 logs at 10 h and recovered to wild-type levels by 14 h. Growth of these two mutants was comparable to wild-type growth in CP-treated and Fe-depleted cocultures (**Figures 7C** and **7E**).

In agreement with prior observations (17, 23), coculture of *S. aureus* with wild-type *P. aeruginosa* in metal-replete CDM led to near-complete killing of *S. aureus* by the 10–12 h timepoints (**Figures 7B** and **S11B**), and Fe-depleted cocultures showed accelerated *S. aureus* killing relative to metal-replete cocultures (**Figures 7D** and **S11B**). Also consistent with prior work (20, 23, 24), the Δ*pqsA-C* mutant exhibited no significant anti-staphylococcal activity in any of the coculture conditions (**Figures 7** **and S14B**), whereas growth of *S. aureus* in coculture with the Δ*pqsH* mutant resembled growth with wild-type *P. aeruginosa* across all coculture conditions (**Figures 7**, **S11** and **S12**). This latter observation is consistent with earlier work showing that PQS alone does not possess significant anti-staphylococcal activity (23, 24).

Notably, growth of *S. aureus* in coculture with the *P. aeruginosa* Δ*pqsL* mutant was distinct from growth with either the wild-type, Δ*pqsH*, or Δ*pqsA-C* strains. In metal-replete CDM, the Δ*pqsL* / JE2 strain combination resulted in a small increase in *S. aureus* viability within the 6–8 h timeframe, and viability was reduced to what was observed in wild-type and Δ*pqsH* cocultures by 14 h (**Figure 7B**). In Fe-depleted CDM, the anti-staphylococcal activity of the Δ*pqsL* mutant was more clearly reduced as compared to wild-type, though not to the levels observed in Δ*pqsA-C* cocultures (**Figure 7D**). These results uncovered that AQNOs are not the only contributors to anti-staphylococcal activity during coculture, which is somewhat surprising given prior work showing that Δ*pqsL* mutants exhibit negligible anti-staphylococcal activity (20, 24).

We then examined the impact of lost AQ production on CP– and ΔΔ-dependent changes in *S. aureus* viability during coculture. As previously observed (17), treatment with either CP or ΔΔ increased *S. aureus* viability during wild-type coculture at the 8–14 h timepoints (**Figures 7F**, **7H** and **S11B**). Intriguingly, coculture with the Δ*pqsL* mutant resulted in significantly enhanced *S. aureus* survival relative to coculture with wild-type *P. aeruginosa* in CP– and ΔΔ-treated cocultures (**Figures 7F** and **7H**). We also noted that loss of AQNO production by the Δ*pqsL* mutant did not rescue *S. aureus* viability in ΔΔ-treated cocultures to the same extent as the loss of all AQ production with the Δ*pqsA-C* mutant (**Figures 7F** and **7H**). Collectively, our findings suggest that the protective effect of CP and ΔΔ on *S. aureus* during coculture with *P. aeruginosa* is primarily attributable to attenuated AQ production in the presence of CP (**Figure 1B, Q2**).

### AQ production contributes to CP– and ΔΔ-dependent changes in cell envelope remodeling responses during coculture

We next examined whether altered pseudomonal AQ levels are responsible for CP– and ΔΔ-dependent changes in cell envelope remodeling gene expression, addressing a key question in our current working model (**Figure 1B**, **Q3**) (18, 100–102). We cocultured *S. aureus* with the *P. aeruginosa* wild-type and above mentioned Δ*pqs* mutants in CP-treated, ΔΔ-treated, and Fe-depleted conditions, and performed real-time PCR to determine the transcriptional responses of genes associated with membrane remodeling in each strain combination. The Δ*pqs* mutants used in this study were constructed from distinct PA14 strains, and our expression data revealed modestly different transcriptional responses between the two parental laboratory strains (**Figures S15A**, **S15C**, **S16A**,**S16C** and **Table S3)**. Thus, gene expression changes for each Δ*pqs* mutant were analyzed relative to the following PA14 parent strains: AO-PA14 for Δ*pqsH*, LD-PA14 for Δ*pqsL* and Δ*pqsA-C*.

For AO-PA14 cocultured with *S. aureus*, real-time PCR showed that CP treatment led to increased expression of membrane remodeling genes associated with lipid A modification (*arnT*) (103), cellular responses to alterations in surface charge (*pmrA*), and the release of eDNA (*cidA*) (51, 104) (**Figure 8**). ΔΔ similarly resulted in the induction of *arnT* and *pmrA*, but not *cidA*, in AO-PA14 cocultures (**Figure 8**). These findings were consistent with transcriptomics and gene expression data presented in this work and in previous studies (13, 17, 18). Expression changes in *arnT* and *pmrA* were largely similar across conditions (**Figure S15**), consistent with *arn* operon expression falling under control of the PmrAB two-component system (56, 57). Overall trends in the expression of membrane remodeling genes for PA14 Δ*pqsH* / *S. aureus* cocultures resembled those of AO-PA14 cocultures (**Figures S15A** and **S15B**), indicating that induction of *P. aeruginosa* cell envelope remodeling responses is not solely dependent on PQS.

**Figure 8.**
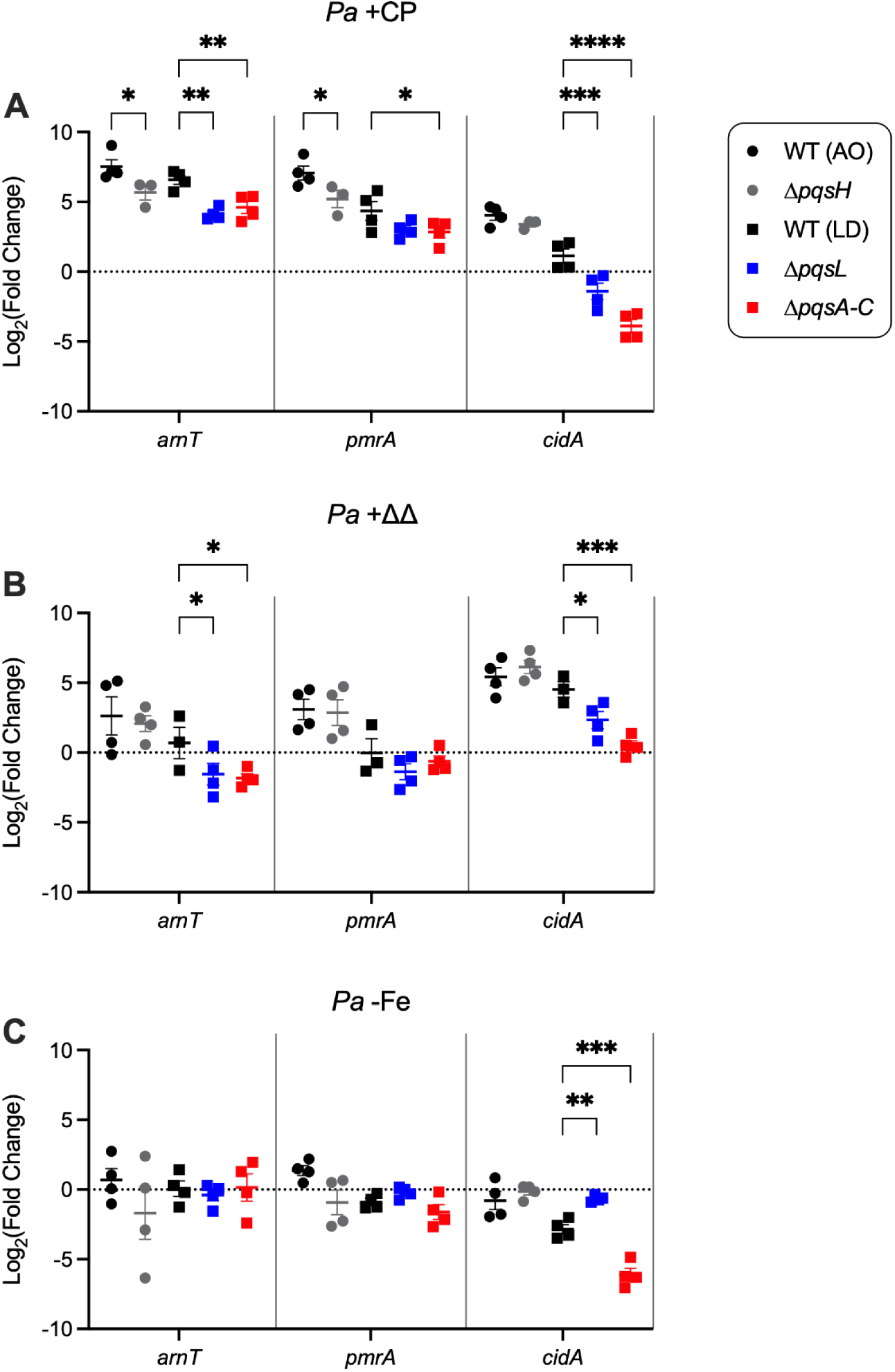
Gene expression changes of *P. aeruginosa* PA14 parent strains and Δ*pqs* mutants in coculture with *S. aureus*. AO WT refers to Oglesby PA14, parent strain of Δ*pqsH* mutant; LD WT refers to Dietrich PA14, parent strain of Δ*pqsL* and Δ*pqsA-C* mutants. Cocultures were grown in CP-treated (**A**), ΔΔ-treated (**B**) or Fe-depleted CDM (**C**) and incubated at 37 °C for 6 h. Transcript levels were normalized to the 16S housekeeping gene for *P. aeruginosa*, and the normalized Log_2_(Fold Change) in gene expression as compared to metal-replete CDM are presented. Statistical significance was calculated by two-way ANOVA relative to parent strain (n ≥ 3, \**P* < 0.05, \*\**P* < 0.01, \*\*\**P* < 0.001, \*\*\*\**P* < 0.0001, error bars represent S.E.). Strainwise comparisons are presented in **Figure S15**.

When considering the *P. aeruginosa* Δ*pqsL* and Δ*pqsA-C* mutant cocultures, we noted that CP treatment of the LD-PA14 strain showed muted induction of *pmrA* as compared to the AO-PA14 strain and no induction of *cidA* (**Figures S15A** and **S15C**). We also observed no induction of the *arnT* and *pmrA* genes in the LD-PA14 cocultures upon ΔΔ treatment (**Figure S15C**). Regardless, CP-dependent changes in *arnT* and *cidA* expression were significantly altered in the Δ*pqsL* / *S. aureus* coculuure as compared to the LD-PA14 coculture, and CP-dependent changes in the expression of all three *P. aeruginosa* membrane remodeling genes were significantly altered in the Δ*pqsA-C* / *S. aureus* coculture as compared to the LD-PA14 / *S. aureus* coculture (**Figure 8A**). The impact of the Δ*pqsL* and Δ*pqsA-C* mutations was more modest in ΔΔ-treated cocultures, with small but statistically significant changes in the induction of *arnT* and *cidA* in these mutant cocultures as compared to LD-PA14 cocultures (**Figure 8B**). These results demonstrated that AQ production is important for the induction of *P. aeruginosa* membrane remodeling responses elicited by the CP protein scaffold during coculture, and that these responses are largely due to the effects of AQNOs with additional AQs also contributing to these gene expression changes.

We also questioned how conditions of Fe depletion would affect the expression of *P. aeruginosa* genes involved in cell envelope remodeling in the Δ*pqs* mutant cocultures (**Figure 8C**), as well as whether the effects of Fe depletion would be similar to or distinct from that of CP treatment (**Figures 8A and 8B**). Consistent with prior work (13, 18), we observed clear differences between the effects of Fe depletion and CP treatment on the expression of cell envelope remodeling genes in wild-type *P. aeruginosa*: specifically, growth in Fe-depleted medium resulted in negligible expression changes for *arnT* and *pmrA* across all strain combinations (**Figure S14A**). Strikingly, growth of the Δ*pqsA-C* mutant in both Fe-depleted and CP-treated cultures resulted in the downregulation of *cidA*, whereas ΔΔ had no effect on *cidA* in these conditions (**Figures 8A-C, S14D,** and **S14E**). These comparisons suggested that changes in *cidA* expression in the absence of AQs are dependent on metal availability, not the activity of the CP protein scaffold.

To probe how CP or ΔΔ treatment affect *S. aureus* cell envelope gene expression in cocultures with the Δ*pqs* mutants, we monitored expression levels for *S. aureus* genes involved in capsular polysaccharide biosynthesis (*cap8F*) (64), competence (*comK*) (73) and cell wall biosynthesis (*sgtB*) (66, 105). We previously reported that expression levels of these genes changed in response to CP treatment (17, 18). Owing to its role in cell wall biosynthesis, *sgtB* has also been shown to be upregulated in response to cell stress (106), providing another readout for *S. aureus* viability across our coculture conditions and Δ*pqs* mutants. For *S. aureus* in coculture with wild-type *P. aeruginosa* (**Figure 9** and **Figure S16A**), overall expression trends in CP– and ΔΔ-treated cocultures aligned well with the cellular responses seen by RNA-seq in this and prior work (18). Specifically, CP treatment led to decreased expression of *cap8F* and *sgtB* by *S. aureus* when cocultured with AO-PA14 (**Figures 9A** and **S16A**). ΔΔ treatment similarly resulted in decreased expression of *sgtB* as well as increased expression of *cap8F* (**Figures 9B** and **S16A**). Coculture of *S. aureus* with the Δ*pqsH* mutant (**Figures 9** and **S16B**) afforded *S. aureus* gene expression trends that resembled those seen in cocultures with the AO-PA14 parent strain, indicating that PQS is not primarily responsible for *S. aureus* cell envelope gene expression changes in response to CP or ΔΔ treatment.

**Figure 9.**
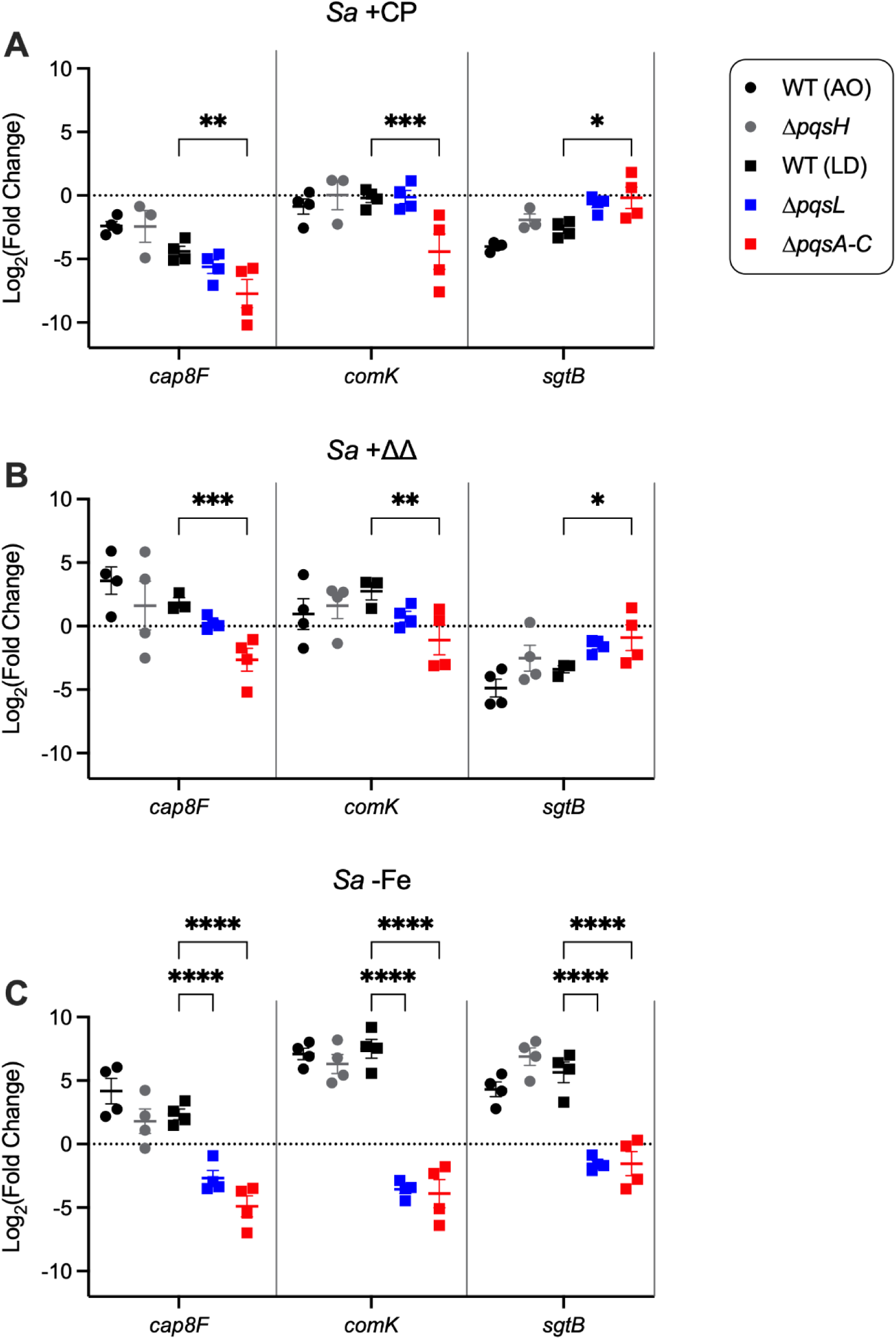
Gene expression changes of *S. aureus* in coculture with *P. aeruginosa* PA14 parent strains and *Δpqs* mutants. AO WT refers to Oglesby PA14, parent strain of Δ*pqsH* mutant; LD WT refers to Dietrich PA14, parent strain of Δ*pqsL* and Δ*pqsA-C* mutants. Cocultures were grown in CP-treated (**A**), ΔΔ-treated (**B**) or Fe-depleted (**C**) CDM and incubated at 37 °C for 6 h. Transcript levels were normalized to the *sigA* housekeeping gene for *S. aureus* and the normalized Log_2_(Fold Change) are presented. Statistical significance was calculated by two-way ANOVA relative to parent strain (n ≥ 3, \**P* < 0.05, \*\**P* < 0.01, \*\*\**P* < 0.001, \*\*\*\**P* < 0.0001, error bars represent S.E.). Strainwise comparisons are presented in **Figure S16**.

The expression profile of *S. aureus* genes during coculture with the LD-PA14 strain was similar to what was observed during coculture with the AO-PA14 strain (**Figures 9**, **S16A** and **S16C**). Compared to a coculture of *S. aureus* with LD-PA14, CP– and ΔΔ-treatment of cocultures with the Δ*pqsL* and Δ*pqsA-C* mutants resulted in reduced expression of *cap8F* and *comK* (**Figures 9A**, **9B** and **S16C-E**). Notably, when comparing across strains, the observed changes in *cap8F* and *comK* expression levels were only statistically significant in the Δ*pqsA-C* mutant (**Figures 9A-B**). In a similar vein, the reduction in *sgtB* expression observed in cocultures with AO-PA14 and LD-PA14 upon CP treatment was not observed in either the Δ*pqsL* or Δ*pqsA-C* cocultures (**Figures 9A**, **S16A**, **S16D** and **S16E**). Moreover, ΔΔ treatment resulted in only a modest reduction of *sgtB* in the Δ*pqsL* mutant cocultures and no reduction in *sgtB* expression in the Δ*pqsA-C* mutant cocultures (**Figures S16D-E**). Combined, these data further indicate that AQNOs are key, but not the sole, drivers of CP-and ΔΔ-dependent changes in the expression of cell envelope modification genes, with AQ QS metabolites also contributing to these gene expression changes.

Lastly, we considered the effects of Fe depletion on *S. aureus* expression of *cap8F*, *comK* and *sgtB* across strain combinations. When *S. aureus* was cocultured with AO-PA14 in Fe-depleted CDM, upregulation of all three genes was observed (**Figure 9C** and **S16A**), which we attributed to increased membrane stress arising from enhanced production of AQNOs by *P. aeruginosa* in Fe-limited conditions (18, 23). Cocultures with the Δ*pqsH* mutant under Fe-depleted conditions afforded *S. aureus* gene expression trends that resembled those seen in cocultures with AO-PA14, again indicating that PQS is not solely responsible for *S. aureus* cell envelope remodeling responses under conditions of Fe depletion. By contrast, the effect of Fe depletion on the expression levels of *cap8F*, *comK* and *sgtB* by *S. aureus* cocultured with the Δ*pqsL* and Δ*pqsA-C* mutants was the reverse of that observed following coculture with the parent strain LD-PA14 (**Figures 9C**, **S16C**, **S16D** and **S16E**). This reversal is likely due to the reliance of *P. aeruginosa* on AQs to kill *S. aureus* during Fe depletion (23); thus, upon loss of AQ production, *S. aureus* experiences a decrease in membrane stress in Fe-depleted cocultures.

Taken together, our real-time PCR studies demonstrate that, within cocultures of *P. aeruginosa* and *S. aureus*, pseudomonal AQNOs play a primary role in inducing transcriptional responses associated with cell envelope remodeling in both species, with secondary contributions from other AQs (**Figure 1B**, **Q3**). Furthermore, our observations show that the CP protein scaffold, without functional metal-binding sites, impacts *P. aeruginosa* and *S. aureus* coculture dynamics by altering AQ production (**Figure 1B, Q1**). Indeed, in the absence of AQs (*i.e.,* in coculture with the Δ*pqsA-C* mutant), the effect of CP treatment on the *S. aureus* membrane remodeling response resembled the effect of Fe depletion (**Figure S16E**), and ΔΔ treatment did not significantly impact *S. aureus* cell viability or membrane remodeling responses (**Figures S14B** and **S16E**). Small differences between expression profiles of *S. aureus* in coculture with Δ*pqsL* and with Δ*pqsA-C* (**Figures S16D** and **S16E**), such as the downregulation of *comK* in CP-treated Δ*pqsA-C* cocultures, indicated that secondary contributions from other AQs beyond the AQNOs can affect *S. aureus* membrane remodeling.

## DISCUSSION

The current study addressed key questions regarding the recently uncovered metal-independent activities of CP that affect interactions between *P. aeruginosa* and *S. aureus* (17, 18). Transcriptomics analyses of ΔΔ-treated cocultures provided robust evidence that the CP protein scaffold elicits gene expression changes indicative of cell envelope modifications for both *P. aeruginosa* and *S. aureus* in coculture. Similar to what was observed with CP treatment (18), quantitative metabolite analyses revealed that ΔΔ treatment resulted in decreased production of the QS metabolite C_4_-HSL and the anti-staphylococcal AQ HQNO by *P. aeruginosa* during coculture. This work further demonstrated that AQs are central to CP– and ΔΔ-dependent changes in gene expression related to cell envelope modifications by both pathogens. Taken together, these observations allowed us to advance the working model describing the consequences of CP on *P. aeruginosa* / *S. aureus* interspecies dynamics (**Figure 10**). We integrated our new findings that CP modulates interspecies dynamics by attenuating AQ production in a metal-independent manner, highlighting how the activities of CP in innate immunity extend beyond nutrient metal sequestration (**Figure 10**). This work also redefined the role of specific AQs in interactions between *P. aeruginosa* and *S. aureus* in the absence and presence of CP. Whereas AQNOs have long been considered the primary or even sole drivers of interactions between these species (20, 24, 107), our current results convincingly show that additional AQs are critical players in *P. aeruginosa / S. aureus* coculture dynamics. These results provide compelling evidence that the metal sequestering properties of CP and the protein scaffold exert intertwined effects on *P. aeruginosa* / *S. aureus* coculture dynamics.

**Figure 10.**
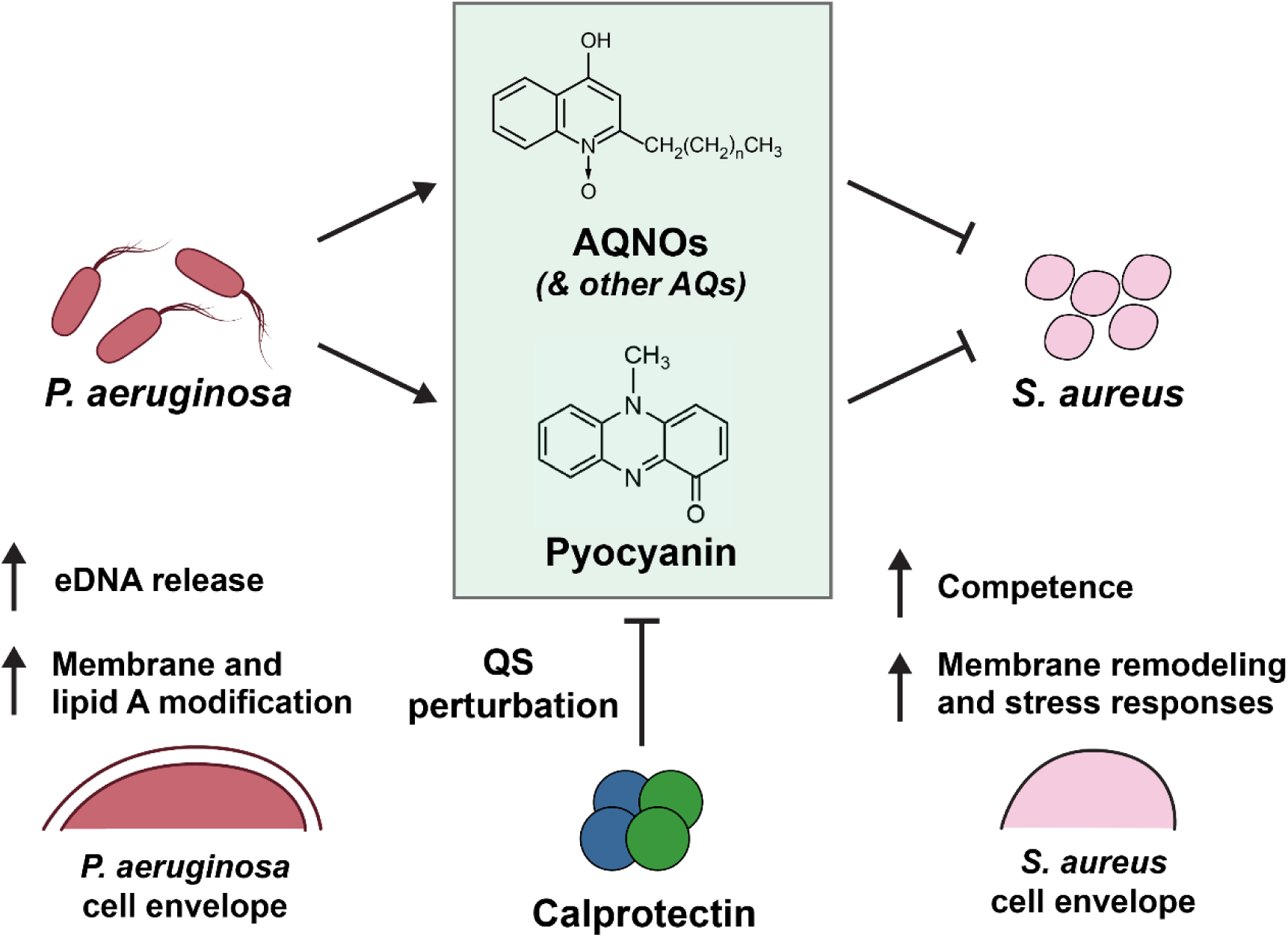
Revised working model depicting how the calprotectin protein scaffold contributes to *P. aeruginosa* / *S. aureus* interspecies dynamics. The CP protein scaffold perturbs pseudomonal QS and in turn decreases the production of anti-staphylococcal factors and promotes *S. aureus* survival in coculture. Pseudomonal AQs elicit gene expression changes indicative of cell envelope remodeling responses from both *P. aeruginosa* and *S. aureus* in coculture. Pyocyanin is included in the model based on prior work (14, 17, 18).

An outstanding question that emerges from our prior and current data is how the CP protein scaffold affects bacterial physiology. Within the context of nutritional immunity, CP activity has long been thought to be contact-independent (2, 6, 7). Nevertheless, some investigations have probed contact dependence, providing evidence for physical interactions between CP and various organisms (108–110). Given the CP– and ΔΔ-dependent changes to the expression of genes involved in modifications to the cell envelopes of both *P. aeruginosa* and *S. aureus*, it will be informative to determine whether physical interactions of the protein with one or both of these pathogens occur and contribute to its ability to perturb QS and AQ production. Along these lines, CP was shown to form a “mesh-like” structure that contacted and encased *P. aeruginosa* and *S. aureus* in statically-grown biofilm cocultures (111). This observation, combined with our current work, raises the question of whether the mesh-like structure has the capacity to affect signal transduction responses in either pathogen, leading to the observed changes in gene expression, metabolite production, and coculture dynamics. In *P. aeruginosa*, the formation, release, and fusion of outer membrane vesicles (OMVs) are integral to QS (97, 112–114), and one possibility worth consideration is whether physical interactions between the protein scaffold and *P. aeruginosa* affect OMV assembly and transport.

Our prior working model posited that altered QS and AQ production by *P. aeruginosa* in the presence of CP was due to altered chorismate flux (18). Chorismate is a biosynthetic precursor for the AQs, as well as PCH and phenazines, and our transcriptomics study of CP-treated cocultures revealed changes in the expression of the respective *P. aeruginosa* genes involved in biosynthesis of these metabolites, consistent with changes in metabolite levels detected in coculture supernatants (17). In contrast, our current transcriptomics analysis of ΔΔ-treated cocultures showed negligible changes in the expression of these genes. This outcome was somewhat surprising given our prior finding that ΔΔ treatment decreased production of the phenazine pyocyanin (PYO) (17) and the metabolite analyses presented in this work. It is possible that ΔΔ treatment affected expression of chorismate metabolic genes, but that these changes were not evident at the timepoint selected for this dataset. It is also possible that decreased PYO production in the presence of CP and ΔΔ is due to decreased C_4_-HSL production (**Figure 5C**), as C_4_-HSL coactivates the expression of genes for PYO production with RhlR (27, 84). Investigation of these distinct yet not mutually exclusive possibilities warrants future consideration.

The observation that the CP protein scaffold impacts the expression of metal homeostasis genes for *S. aureus* in coculture and for *P. aeruginosa* in both mono– and cocultures in this study was also surprising. In the case of *P. aeruginosa*, multiple genes associated with Fe acquisition machinery were downregulated following ΔΔ treatment (**Figures S3A** and **S4**), and elucidating the origins of this response is an intriguing topic for future investigation. By contrast, *S. aureus* upregulated the expression of genes associated with Fe and Zn acquisition during coculture as a consequence of ΔΔ treatment (**Figure S5**). This result also requires investigation and could be due to increased *S. aureus* fitness in the presence of ΔΔ resulting from decreased AQ production by *P. aeruginosa* (see Supplementary Discussion).

Although CP and ΔΔ promoted the growth of *S. aureus* in coculture with *P. aeruginosa* to a similar extent (17), a comparison of our dual-species transcriptomic datasets uncovered clear differences in the transcriptional responses of *P. aeruginosa* and *S. aureus* in coculture to CP and ΔΔ treatment. We observed much smaller effects of ΔΔ treatment on the transcriptomes of both pathogens during coculture when compared to previously observed effects of CP treatment. Moreover, we found that many of the changes elicited by ΔΔ from each organism were specific to coculture. Changes in gene expression and coculture dynamics upon ΔΔ treatment further appeared to be dependent on *P. aeruginosa* AQ production. These findings suggested that the combination of CP-mediated metal sequestration and the metal-independent activities of the CP protein scaffold were required to induce gene expression changes in a subset of genes, and that these effects were dependent on coculture interactions between these species. Collectively, this work provides compelling support for the notion that CP simultaneously exerts canonical metal-dependent and non-canonical metal-independent effects when contributing to host defense and modulating pathogen–pathogen interactions.

## EXPERIMENTAL PROCEDURES

### General materials and methods

#### Solutions, buffers and metal stocks

All chemicals and reagents were commercially obtained and used as received. Solutions, buffers and metal stocks were prepared with Milli-Q water (18.2 MΩ·cm, Milli-Q Academic) and filtered (0.2 μm) before use as previously described (17, 18).

#### Protein overexpression, purification and handling

CP and ΔΔ were overexpressed and purified as previously described (7, 17, 29). Protein stocks were thawed and prepared for use in microbiology as previously described (17). Briefly, protein aliquots were thawed and buffer exchanged against ice-cold 20 mM Tris (VWR, 97061), 100 mM NaCl pH 7.5 (AMA buffer) using an Amicon Ultra-0.5 centrifugal spin-filter (MWCO 10 kDa, UFC5010BK). The final concentration of protein was determined using absorbance measurements at 280 nm (17).

#### Statistical testing

Unless otherwise stated, significance values were calculated as compared to the untreated (metal-replete condition) using Welch’s unpaired *t*-test.

### Microbiology

#### Growth media

Luria-Bertani (LB, Miller) medium (BD Difco), *Pseudomonas* Isolation Agar (Sigma, BD Difco) and Baird-Parker medium (Sigma, BD Difco) were commercially obtained, dissolved into Milli-Q water and prepared according to manufacturer recommendations. Sterilized Baird-Parker medium was supplemented with egg yolk tellurite emulsion (Sigma, BD Difco). Chemically defined medium (CDM) was prepared as previously reported (8, 17). Unsupplemented CDM was prepared from trace metals basis reagents, and metal-replete CDM was prepared by addition of 1 mM Ca(II), 0.3 µM Mn(II), 5 µM Fe(II), 0.1 µM Ni(II), 0.1 µM Cu(II) and 6 µM Zn(II) to unsupplemented CDM immediately prior to usage. Fe-depleted CDM was prepared by omitting Fe from the above supplementation step.

#### General methods for bacterial culture

General microbiology methods, including strain handling, colony-forming unit (CFU) counting and turbidity measurements, were performed as previously reported (17). For bacterial cultures involving protein treatment, 20 µM of protein or the equivalent volume of AMA buffer was added. Cocultures of *P. aeruginosa* and *S. aureus* were inoculated using a starting OD_600_ of 0.005 and 0.02 respectively (log(CFU/ml) = 6.8– 7.0 and 7.2–7.4 respectively), and unless otherwise stated, monocultures of each species were inoculated using the same starting OD_600_ values. The complete list of bacterial strains used is tabulated in **Table S3**.

#### RNA extraction, workup, cDNA synthesis and real-time PCR

RNA extraction and workup from bacterial cultures, and the subsequent steps to prepare cDNA for real-time PCR analysis were conducted as previously reported (17, 18). In short, bacterial cell pellets were isolated from cultures at the 6 h timepoint, lysed, and RNA was extracted using phenol/chloroform/isoamyl alcohol (Sigma, 77617). RNA was then purified using the Qiagen RNeasy kit with two rounds of DNAseI treatment. The purified RNA was then precipitated overnight in ice-cold 100% ethanol containing 0.3M NaOAc. The precipitated RNA was washed and the residual ethanol was allowed to evaporate at ambient temperature, following which the RNA pellet was resuspended in nuclease-free H_2_O (VWR, E476) and the final concentration of isolated RNA was determined using a Synergy HT microplate reader (Biotek). cDNA synthesis was carried out using the Protoscript II First Stand cDNA Synthesis kit (NEB) according to manufacturer instructions. For validation of RNA-seq gene expression change, real-time PCR standard curves were performed as previously reported for each gene to determine the amount of input RNA used for cDNA synthesis (17). The following amounts of input RNA were used for each gene: *lasA*: 10 ng; *pvdS*: 300ng; *flp*: 300 ng. For studies of cell envelope remodeling responses in the cocultures using *P. aeruginosa* Δ*pqs* genetic mutants, standard curves were produced for each primer set by analyzing cDNA from serial dilutions of RNA, ensuring the C_t_ range of the standard curve encapsulated collected C_t_ values across all conditions. Standard curves were used to determine the relative amount of RNA in each sample as previously described (115), and relative RNA levels were normalized according to *16S* and *sigA* RNA levels. 300 ng input RNA was used for all genes under all conditions. The full list of primers used for real-time PCR is listed in **Table S4**, and the input RNA ranges that afforded robust standard curves for each gene of interest are listed in **Table S5** and **Table S6**.

#### RNA-seq and bioinformatics analyses

Sample preparation for RNA-seq and downstream bioinformatics processing were conducted as previously reported (18). Briefly, isolated RNA samples were submitted to the MIT BioMicro Center for quality control, library preparation and subsequently RNA-seq on an Ilumina NextSeq 500 instrument (75 nt chemistry). The obtained raw reads were aligned to the combined reference genomes available for *P. aeruginosa* PA14 and *S. aureus* JE2 on the NCBI RefSeq database (18). The aligned reads were then quantified using Feature Aggregate Depth Utility (tool for prokaryotic read sample quantification) (116). With the feature counts in hand, differential expression analysis was performed using DESeq2, and overexpression analysis was performed using clusterProfiler (18, 117–119). Principal component analysis was also performed on the obtained libraries to check for technical artifacts and errors (**Figure S17** and **Figure S18**).

#### Triple-quadrupole mass spectrometry

Triple-quadrupole mass spectrometry (MS) was performed on an Ultivo triple-quadrupole instrument fitted to an Agilent 1260 Infinity System following previously established methodology (18). In brief, the instrument was run in multiple reaction monitoring mode for simultaneous detection and quantification of AQs and homoserine lactones extracted from cell-free culture supernatants. Analyte standards were prepared from commercially available sources and used to prepare standard curves for determining the dynamic range of detection for each analyte (**Table S7**). The optimal run parameters were obtained using MassHunter Optimizer (Agilent) software (18). Cell-free supernatants were obtained by centrifugation, spiked with internal standard, and extracted with ethyl acetate containing 0.02% v/v acetic acid. Following phase separation, the organic layers were combined and the solvent was evaporated. The extracted analytes were redissolved in ice-cold methanol and further processed for mass spectrometric analysis.

## DATA AVAILABILITY

RNA-seq raw reads were deposited in the NCBI Sequence Read Archive under the BioProject accession number XXXXXXX. The full list of differentially expressed *P. aeruginosa* and *S. aureus* genes are presented in the accompanying Supplemental Excel file.

## ACKNOWLEDGEMENTS

The authors would like to acknowledge Dr. Mohanraja Kumar (MIT DCIF) for technical assistance and expertise regarding MS. The authors would also like to thank Prof. Dianne Newman (Caltech) and Prof. Lars Dietrich (Columbia) for providing bacterial strains used in this work.

## FUNDING AND ADDITIONAL INFORMATION

This work was supported by the NIH (R01 GM118695 to E.M.N. and R01 GM126376 to E.M.N. and A.G.O.). W.H.L. is a recipient of the A*STAR National Science Scholarship (BS-PhD). The triple quadrupole MS instrument is housed and maintained in the MIT DCIF (HHMI). The real-time PCR and RNA-seq instruments are housed and maintained in the MIT BioMicro Center (NIH P30-ES002109).

## DISCLAIMER

The content is solely the responsibility of the authors and does not necessarily represent the official views of the National Institutes of Health.

